# N1 component of event-related potential evoked by simple composite figures in the lateral occipital cortex

**DOI:** 10.1101/345934

**Authors:** Hiroyuki Uchiyama, Shoichi Iwashita, Takashi Mitsui

## Abstract

Since Sugawara and Morotomi (1991) first reported that illusory contour (IC) figures, or Kanizsa square inducers (KSI), evoked a significant N1 component, the first visually evoked negative component of event-related potentials (ERPs), in the lateral occipital cortex (LO), many ERP studies have confirmed their finding, and these studies showed that Kanizsa-type inducers alone evoked a relatively large N1 in the LO, even when the inducers are oriented to not form an IC (see Murray and Herrmann, 2013). In the present study, we used non-IC simple composite figures composed of several elementary shapes (e.g., triplet squares or quadruplet circles) as visual stimuli, and found that the composite figures evoked a significant N1 comparable to that evoked by KSI in the anterior LO, whereas solitary elementary shapes did not. Thus, it appears that the LO is activated by not only an IC but also the compositeness of figures, which implies that the LO might analyze the geometrical relationships among multiple 2D shapes that compose a single composite figure.

## Introduction

Event-related potentials (ERPs) are a useful method for analyzing various neural processes, including visual processing, primarily because of their much higher temporal resolution than functional magnetic resonance imaging (fMRI) and positron emission tomography (PET) (Luck, 2014). The occipital N1 component is the first negative ERP component evoked by visual stimuli, and it likely reflects the neural operation of an attentive visual discrimination process (Vogel & Luck, 2000). The present study used the N1 component as an index of neural activity for shape analysis in the lateral occipital cortex (LO).

Sugawara and Morotomi first reported that illusory contour (IC) figures, including Kanizsa square inducers (KSI), evoked a significant N1 in the LO (Sugawara & Morotomi, 1991). To date, many studies, including fMRI studies, have investigated the activation in the LO evoked by IC figures (see (M. M. Murray & Herrmann, 2013) for a review). These studies have indicated that Kanizsa-type inducers alone evoke a significant N1 in the LO, even when the inducers are oriented to not form an IC. Moreover, these inducers can evoke a much larger N1 in the LO if they are oriented to form an IC.

Uchiyama and colleagues created a computerized navigation test, the Random Walker Test (RWT), to quantitatively assess left-right orientation ability (Uchiyama, Mitsuishi, & Ohno, 2009). The RWT uses a set of three square graphical user interface (GUI) buttons arranged in a pyramid shape to enable participant directional choice (left, front and right). Uchiyama and colleagues accidentally found that regardless of the presence or absence of left-right orientation, triplet squares arranged in a pyramid shape (pyramidal triplet squares, PTS) visually evoked a large N1 at lateral occipital sites using a conventional 10-20 EEG recording system (unpublished observation). In the present study, we first localized the scalp distribution and the sources of N1 evoked by PTS with a high spatial resolution via high-density ERP recordings using 64 electrodes. In the first experiment, we also presented pictures of human faces as a response benchmark for N1 because faces are widely believed to be the most favored visual stimuli for visual N1 or N170 (Luck, 2014; Rossion & Jacques, 2011). Our experiment showed that the N1 component evoked by PTS was bilaterally localized on the anterior LO (LO2), and the response on the right side was 23% larger than the response on the left side. Faces also evoked a large N1 component as previously reported, but this response was not localized in the anterior LO. In the second experiment, we compared the N1 amplitudes evoked at the right anterior lateral occipital site (PO8) by simple composite figures composed of elementary shapes, such as PTS and quadruplet circles (QC), with the amplitudes evoked by solitary elementary shapes. In the third experiment, we compared the N1 amplitudes evoked by five figures composed of three squares (long rectangle, square and rectangle, L-shape, PTS and oblique triplet squares) at PO8. These comparisons showed that simple composite figures evoked a significant N1 comparable to that evoked by KSI localized in the anterior LO, whereas solitary elementary shapes did not. If shape elements of the composite figures were physically contacted or if partial occlusions of certain element shapes were subjectively perceived, the anterior LO was further activated. Thus, the anterior LO might not analyze the contour or surface of solitary elementary shapes. Rather, it might analyze the geometrical relationships among multiple two-dimensional (2D) shapes that compose a single composite figure.

## Material and methods

The experimental protocols of the present study were in accordance with the ethical guidelines of noninvasive brain research in humans of the Japan Neuroscience Society and approved by the local ethics committee for human research at Kagoshima University Graduate School of Engineering and Science.

We examined 57 right-handed male Japanese university students (19-25 years old, average = 22 years). The HN Handedness Inventory was used to quantitatively identify handedness (Hatta & Nakatsuka, 1976). All participants had normal or corrected-to-normal vision. The visual stimulation was presented using a notebook PC (Dynabook B550, Toshiba, Tokyo, Japan). The presentation of stimuli was controlled using MATLAB 2013B (The MathWorks, Natick, MA) and MATLAB scripts. Trigger signals were sent from the PC to the electroencephalogram (EEG) polygraph (EEG-1200, Nihon Koden, Tokyo, Japan) through a data IO interface (NI-USB6008, National Instruments Japan, Tokyo, Japan).

Scalp potentials were recorded through an array of 64 low-noise active electrodes (actiCAP, Brain Products, München, Germany), and the impedance of the electrodes was maintainde below 10 kΩ. The electrodes were placed according to the 10-10 system (Fig. S1). An additional electrode was placed at AFz as the ground. Electroencephalography was recorded using the connection of Fz and F4 as a reference. The time constant of the polygraph was set at 0.3 s, and the cut-off frequency of the low-pass filter was 60 Hz. The waveform data of the 64 channels were digitized at a 1-kHz sampling rate and stored. Waveform data were digitally filtered between 1 and 40 Hz and were averaged for each type of stimulus using BESA Research 6.0 (BESA GmbH, Gräfelfing, Germany). Next, the waveform data of the Fz channel were subtracted from the waveform data of all channels so that Fz could be regarded as the sole reference. Data from some participants (6 participants in Experiment 1, 4 participants in Experiment 2 and 4 participants in Experiment 3) were omitted from further analysis, because of noise, blink and body movement artifacts and a lack of distinguishable N1 components. Therefore, the data from 22 participants in Experiment 1, 16 participants in Experiment 2 and 15 participants in Experiment 3 were analyzed. The N1 peak amplitudes of the participants evoked by shapes and faces were statistically compared; paired t-tests, ANOVAs and paired t-tests modified for multiple comparisons were performed, using MATLAB, R and RStudio. Grand averages of all participants were used for topographical mappings, via MATLAB, EEGLAB (Delorme & Makeig, 2004) and ERPLAB (Lopez-Calderon & Luck, 2014) and the brain electrical source analysis (BESA).

In Experiment 1, color pictures of PTS (Fig. 1a) and human faces were presented to the participants. Pictures of faces were selected from the FEI Face Database (Thomaz & Giraldi, 2010; http://fei.edu.br/~cer/facedatabase.html). The face pictures were 72 mm in height by 97 mm in width on the monitor, and the size of the face area was 40-47 mm by 30-35 mm. The dimensions of the TS ranged from 40 to 60 mm on both sides. The participants viewed the monitor at a distance of 40 cm; therefore, the size of the stimulus ranged from approximately 4° to 9° in visual angle. The luminosity of the display was 0.7-22.9 cd/m^2^. The PTS colors were red, green or blue, and the PTS orientations were upward, downward, leftward or rightward. Thus, 12 TS patterns in various combinations of color and orientation were equally and randomly presented. After the presentation of a fixation point for 0.5-1.0 s or 1.0-1.6 s, a face or PTS image was randomly presented for 1 s. The participants subsequently expressed the color of the TS or the sex of the individual using the GUI buttons via a mouse. Each participant viewed 100 different faces (males/females, 50 each) and 100 PTS.

In Experiment 2, multielement figures and solitary elementary shapes were presented to the participants. The multielement figures included PTS, unconnected triplet squares (UTS; Fig. 1b), QC (Fig. 1c), and KSI (Fig. 1d), and the solitary shapes included a solitary square (SS; Fig. 1e) and a solitary circle (SC; Fig. 1f). QC consisted of four 20-mm-diameter circles and circles that were arranged rectangularly and separated by 10 mm. The top square of the UTS was separated from the base squares by 20 mm. KSI consisted of four 270° major sectors, and each sector was oriented to form an IC with neighboring sectors. The side length of the SS was 35 mm so that its area matched the total area of three 20-mm-by-20-mm squares. The diameter of the SC was 49 mm. As in Experiment 1, the participants determined the color of the presented shapes (green, blue and black). Each of the six types of shapes was presented 60 times for each participant.

**Fig. 1.**
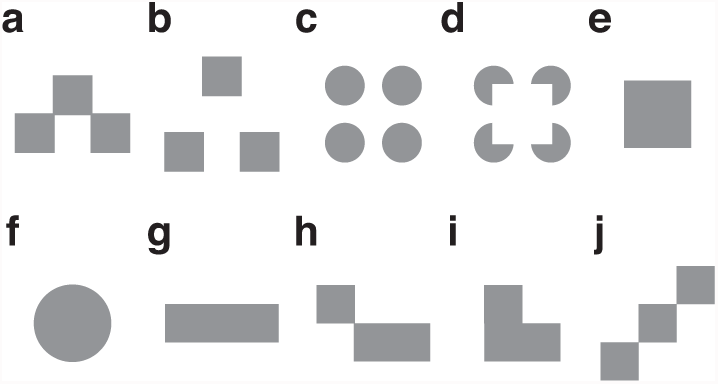
Shapes presented in the present study: pyramidal triplet squares (PTS; a), unconnected triplet squares (UTS; b), quadruplet circles (QC; c), Kanizsa square inducers (KSI; d), solitary square (SS; e), solitary circle (SC; f), long rectangle (LR, g), square and rectangle (SR, h), L-shape (LS, i) and oblique triplet squares (OTS, j). Color versions of the shapes (a-f) were presented in the actual sessions of Experiments 1 and 2, and black versions of the shapes (a, g-i) were presented in Experiment 3. (a, b, h and i) Four rotated versions were equally presented. (g, j) Two rotated versions were equally presented.

In Experiment 3, five figures composed of three squares were presented to the participants. The figures included PTS, long rectangle (LR, Fig. 1g), square and rectangle (SR, Fig. 1h), L-shape (LS, Fig. 1i) and oblique triplet squares (OTS, Fig. 1j). Black pictures of these figures were presented on a gray background for 0.3 s. The participants determined the number of figure elements; the number of elements of LR and LS, SR and PTS and OTS are one, two and three, respectively.

## Results

### Experiment 1 (PTS *vs.* faces)

We presented pictures of PTS (Fig. 1a) and human faces to the participants in a random order and simultaneously recorded ERPs using 64 electrodes based on the 10-10 system (Fig. S1). Fig. 2 shows the grand-averaged (n = 22) ERPs of 14 posterior sites evoked by PTS and faces. Both PTS and faces evoked significant P1, N1 and P2 components at the occipital, posterior parietal, and posterior temporal sites. However, faces evoked much larger P1 than PTS did, particularly at the medial occipital sites (O1, Oz and O2).

**Fig. 2.**
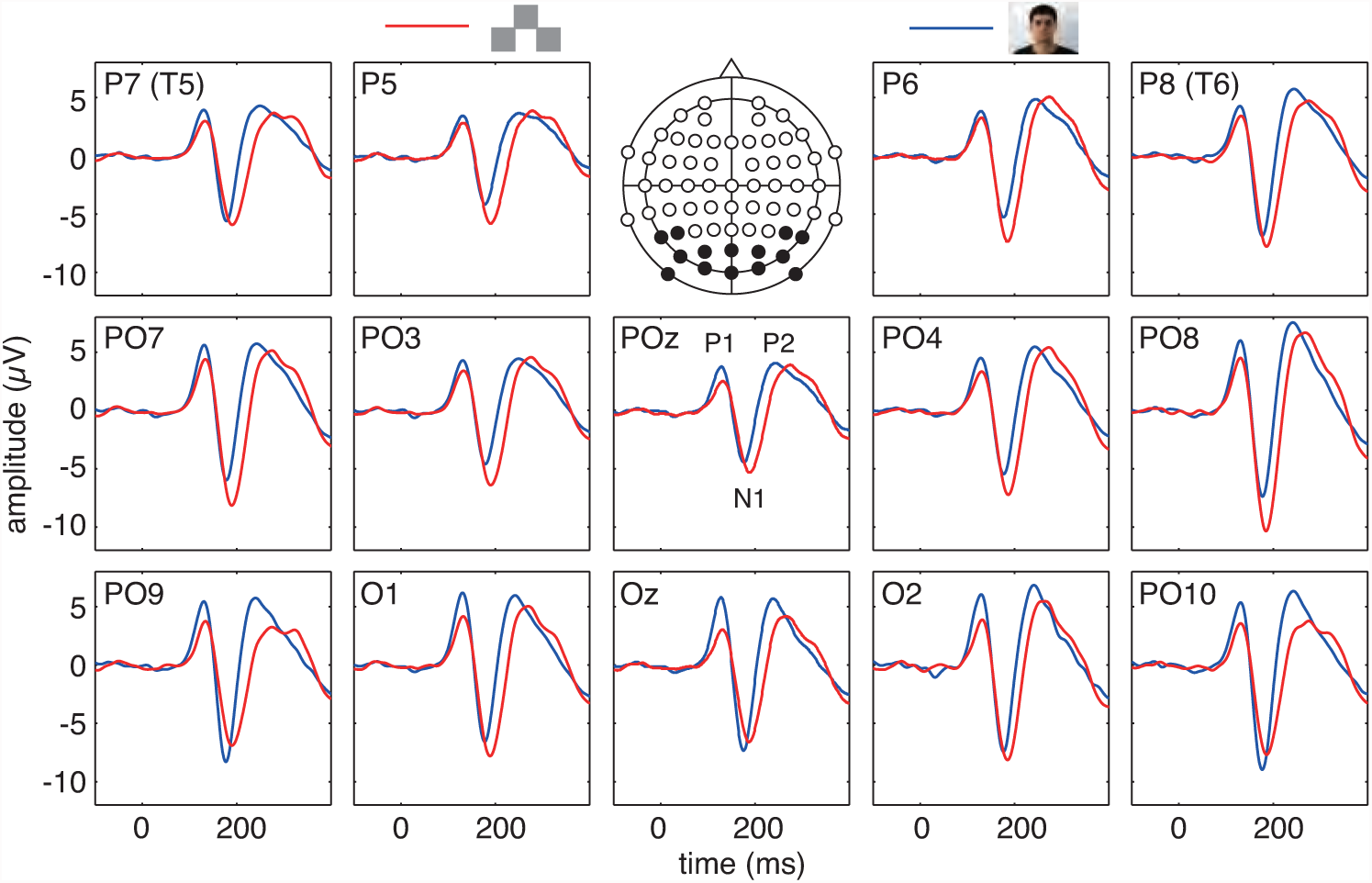
Grand-averaged (n = 22) ERPs evoked by pyramidal triplet squares (PTS; red line) and faces (blue line). Only the ERPs recorded at the 14 posterior sites, indicated in the inset of the top row, are shown here.

The maximal N1 evoked by PTS for each participant was recorded at PO8 (number of participants: 12), PO7 (3), PO4 (2), PO10 (2), O1 (2) and O2 (1), and the maximal N1 evoked by faces for each participant was recorded at PO10 (6), PO9 (4), O2 (4), PO8 (3), TP9 (2), TP10 (1), P7 (1) and O1 (1). Thus, the maximal N1 evoked by PTS across individuals was much more localized at the right anterior lateral occipital sites than the maximal N1 evoked by faces. The maximal N1 amplitude evoked by PTS for each participant ranged from –5.2 to –23.2 µV, and its average was – 11.7 ± 4.9 (mean ± SD) µV. The maximal amplitude of N1 evoked by faces for each participant ranged from –3.8 to –22.5 µV, and its average was –11.6 ± 5.4 µV. These averages are approximately identical (average of the difference of each participant = 0.1 ± 4.7 µV; paired t-test, p = 0.932). Thus, we confirmed that PTS evoked a significant N1, which is comparable to the N1 evoked by faces. The average peak latency of N1 evoked by PTS was 187 ± 9 ms and that evoked by faces was 178 ± 11 ms. The largest grand-averaged N1 evoked by PTS was at PO8 (–9.9 µV, 188 ms) and that evoked by faces was recorded at PO10 (–9.0 µV, 177 ms). The average individual peak data for N1 evoked by PTS at PO8 was –11.2 ± 5.1 µV (186 ± 9 ms) and that evoked by faces at PO10 was –10.2 ± 5.2 µV (177 ± 8 ms). At PO8, the average amplitude of N1 evoked by PTS was 24% larger than the average amplitude evoked by faces (faces, –9.0 ± 5.6 µV, 179 ± 10 ms; paired t-test, p < 0.05).

The scalp topography of the grand-averaged (n = 22) N1 evoked by PTS showed a significantly localized pattern, with prominent peaks at the anterior lateral occipital sites of both sides (Fig. 3) and a particular prominence on the right side (significant around PO8; Fig. 3b). The scalp topography of the grand-averaged (n = 22) N1 evoked by faces in the present study was similar to the results reported by previous studies (Rossion & Jacques, 2011), and the higher potential zone was distributed broadly across the lower and posterior occipital sites (Figs. 3b and 4). Thus, the N1 topographies evoked by PTS and faces differed, indicating that the cortical areas activated by these stimuli are different.

**Fig. 3.**
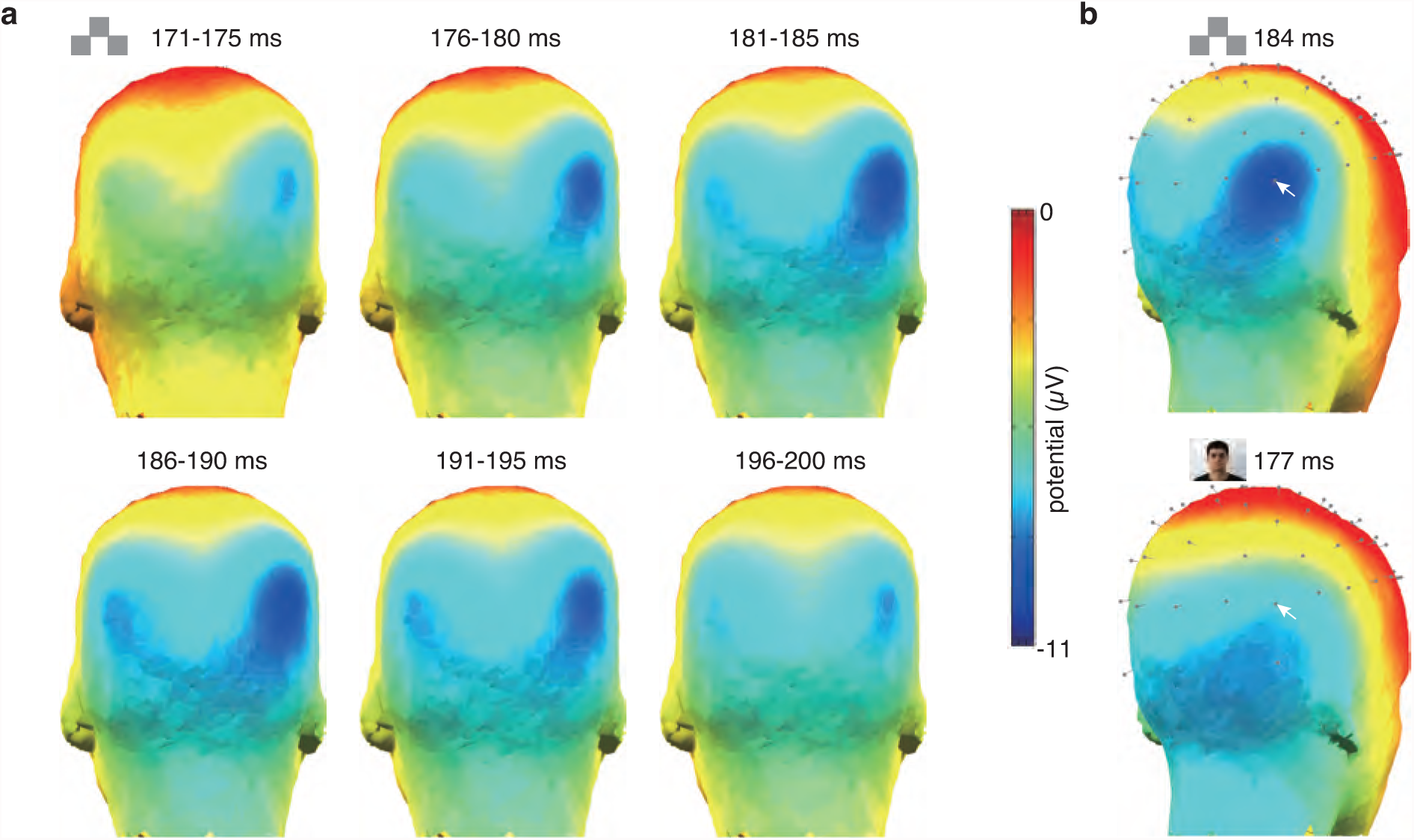
a) Chronological series of scalp topography of the N1 evoked by pyramidal triplet squares (PTS) from 171-175 ms to 196-200 ms. The head is viewed from the back. b) Comparison of N1 scalp topographies evoked by pyramidal triplet squares (PTS; top) and faces (bottom) at their peak time. The heads are rotated so that the right lateral occipital sites are at the center. Gray pins indicate electrode sites, and white arrows indicate the location of PO8.

**Fig. 4.**
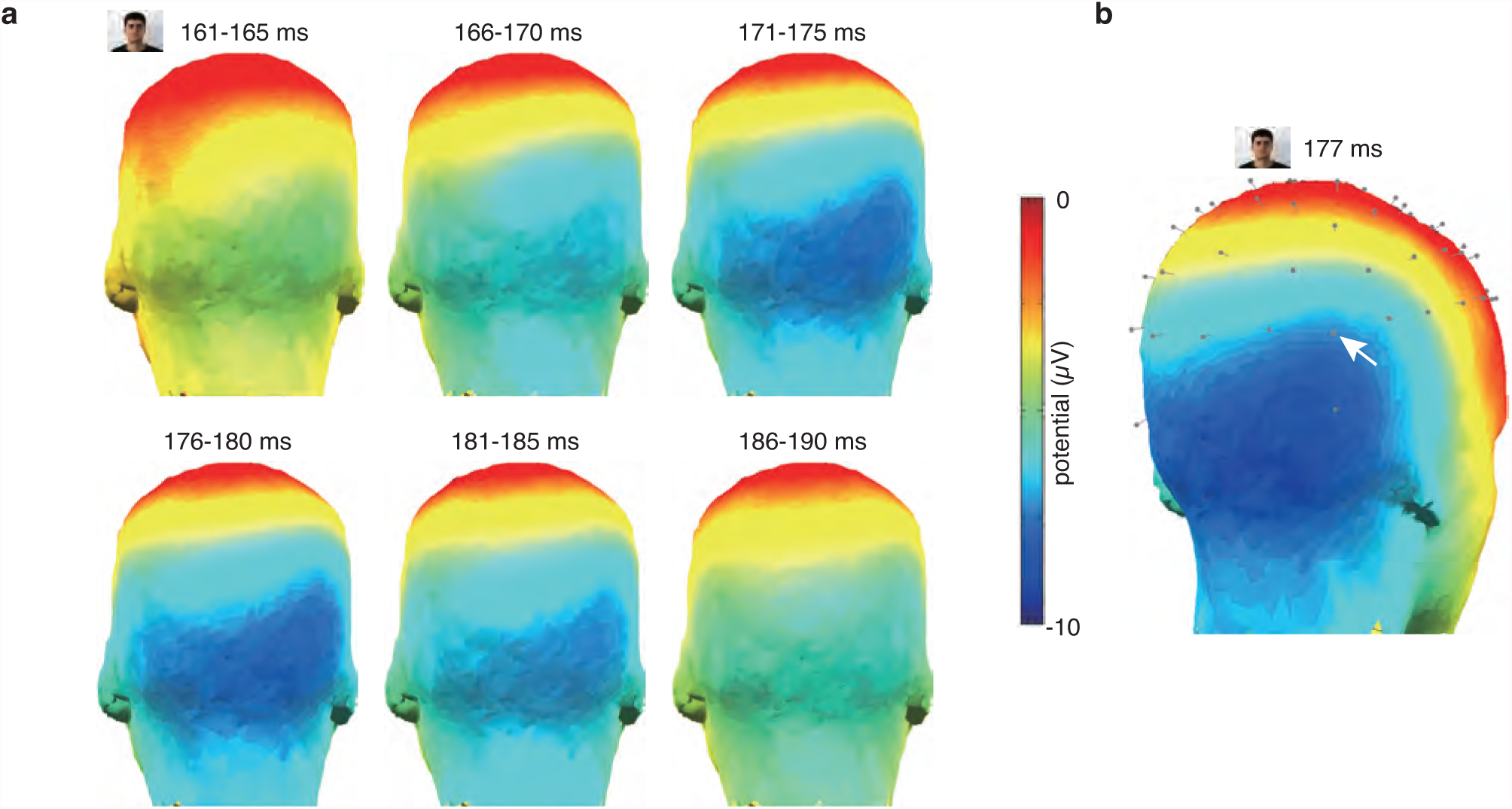
N1 scalp topography evoked by faces. (a) Chronological series from 161-165 ms to 186-190 ms. Heads are viewed from the back. (b) Topography at the peak time (177 ms). The head is rotated so that the right lateral occipital sites are at the center. The white arrow indicates the location of PO8.

The BESA of N1 (time range = 151–220 ms) evoked by PTS revealed that two estimated equivalent current dipoles were located bilaterally in the anterior LO (Fig. S2), and the Talairach coordinates (x, y, z) of the right and left dipoles were (36, –72, –2) and (–30, –67, –5), respectively. Judging from the Talairach coordinates of LO2 by Larsson and Heeger (Larsson & Heeger, 2006), the right dipole was located at the anteromedial edge of the range of the right LO2 ([32, 46], [–89, – 72], [–13, 12]), and the left dipole was closely adjacent anteromedially to the range of the left LO2 ([–43, –31], [–92, –67], [–13, 11]).

The N1 evoked by PTS showed a right-side dominance (Figs. 2 and 3). The average individual N1 peak evoked by PTS at PO8 was 23% larger than that at the corresponding PO7 (–9.1 ± 4.4 µV) in the left lateral occipital region, and the peak latency of N1 evoked by PTS at PO8 was 4 ms faster than the peak latency at PO7 (190 ± 10 ms). A chronological series of the topography maps indicated that the activation evoked by PTS at the right lateral occipital sites started approximately 10 ms earlier than that at the left lateral occipital sites, and the activities of both sides diminished over the same time range (Fig. 3a).

### Experiment 2 (composite figures composed of elementary shapes *vs.* solitary elementary shapes)

We presented four types of multielement figures and two types of solitary shapes to the same participants and compared the ERPs evoked by various types of shapes: multielement shapes (PTS, UTS, QC and KSI; Fig. 1a-d) and solitary shapes (SS and SC; Fig. 1e, f). All shapes evoked distinguishable N1 components at PO8 across 16 participants, but the N1 amplitudes for the different shapes were significantly different (ANOVA, p < 2e–12, Table 1).

**Table 1.**
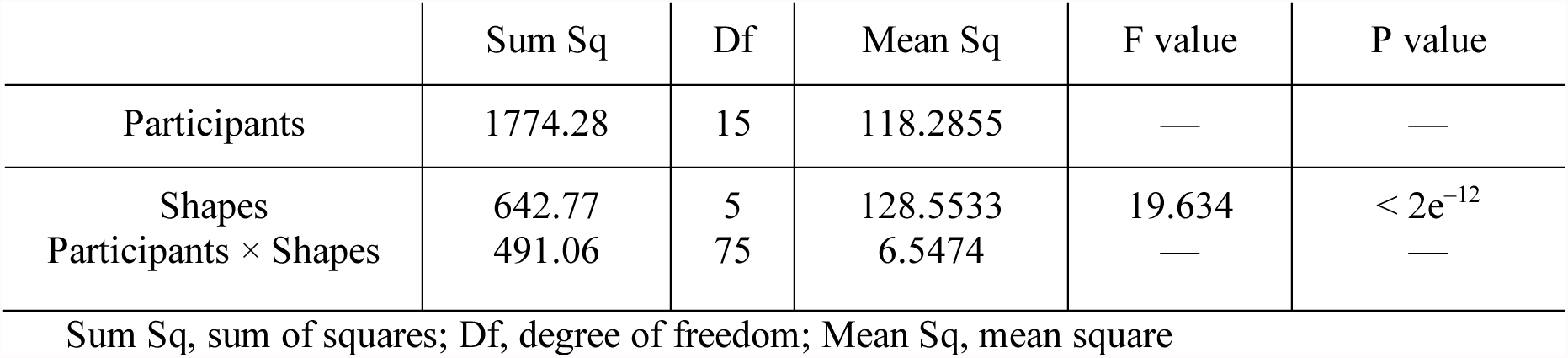
ANOVA of N1 amplitudes evoked by shapes in Experiment 2.

|  | Sum Sq | Df | Mean Sq | F value | P value |
| --- | --- | --- | --- | --- | --- |
| Participants | 1774.28 | 15 | 118.2855 | — | — |
| Shapes | 642.77 | 5 | 128.5533 | 19.634 | $< 2e^{-12}$ |
| Participants × Shapes | 491.06 | 75 | 6.5474 | — | — |

Sum Sq, sum of squares; Df, degree of freedom; Mean Sq, mean square

As in Experiment 1, PTS evoked a large N1 at the lateral occipital sites (at PO8, –12.8 ± 1.3 µV, 192 ± 2 ms, n = 16; Figs. 5 and 6). QC also evoked a large N1 at the lateral occipital sites, and the largest N1 was recorded at PO8 (–9.5 ± 1.4 µV, 185 ± 2 ms; Figs. 6 and 7). The N1 amplitude at PO8 evoked by QC was 26% smaller than that evoked by PTS (paired t-test modified for multiple comparisons, p = 0.0001, Table 2; Fig. 6). Except for peak amplitude and latency, the scalp topography of N1 evoked by QC showed a similar pattern to that evoked by PTS (Fig. 8), indicating that both PTS and QC activate the same cortical area, the anterior LO.

**Fig. 5.**
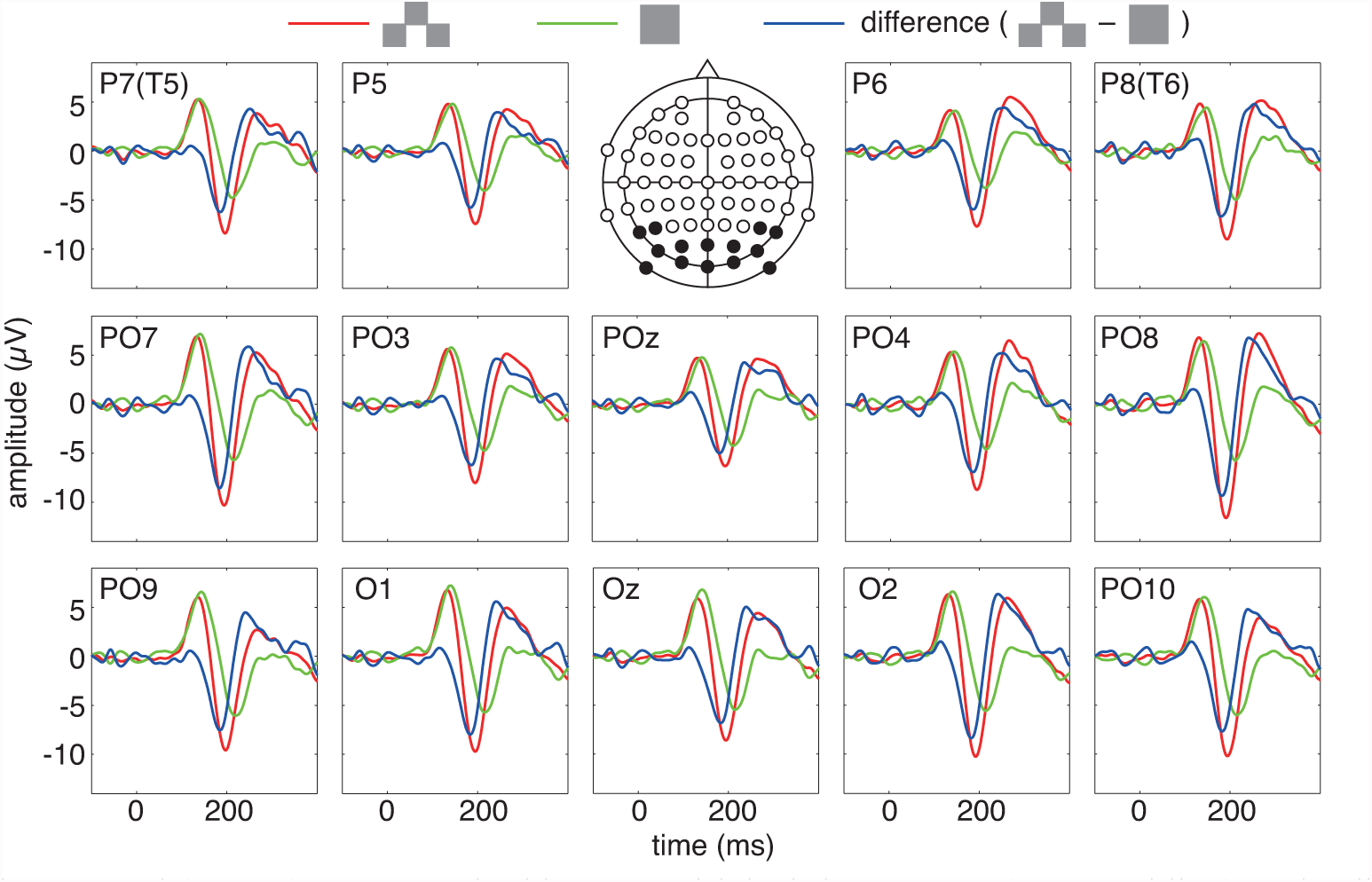
Grand-averaged (n = 16) ERPs evoked by pyramidal triplet squares (PTS; red line) and solitary square (SS; green line) and the difference waves (PTS – SS; blue line).

**Fig. 6.**
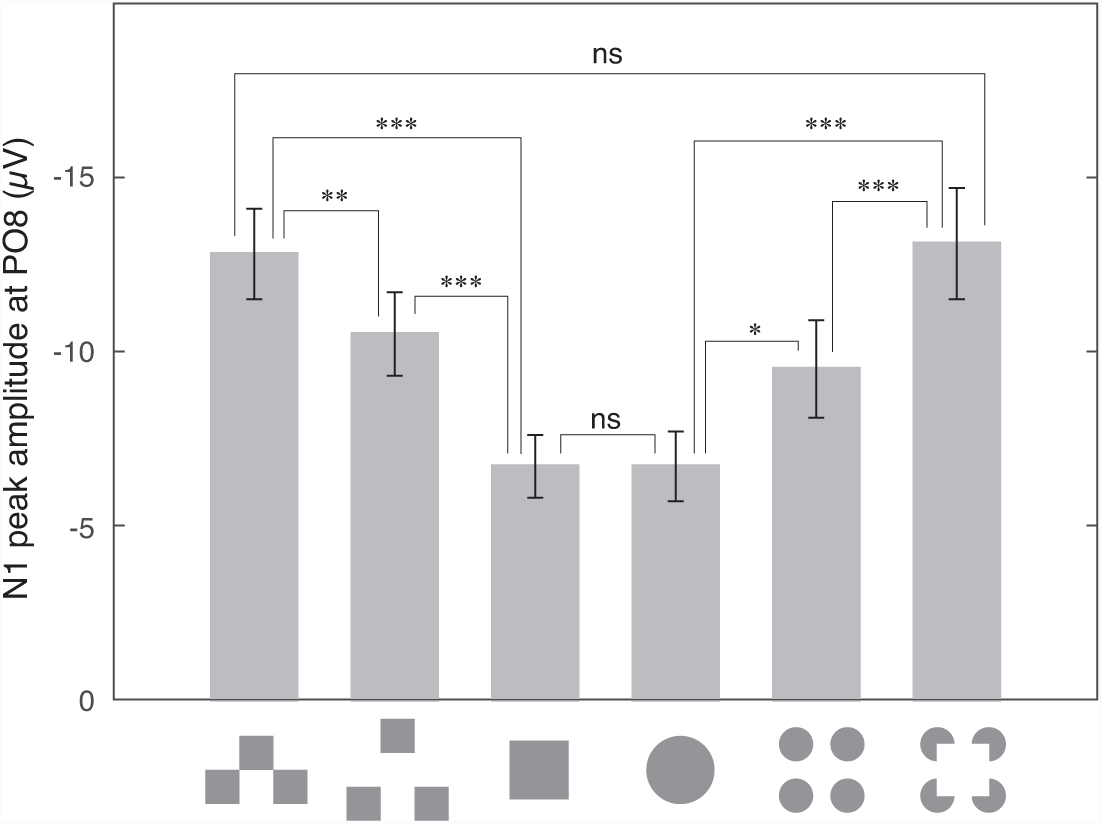
Averaged (n = 16) N1 peak amplitudes evoked by pyramidal triplet squares (PTS), unconnected triplet squares (UTS), solitary square (SS), solitary circle (SC), quadruplet circles (QC) and Kanizsa square inducers (KSI) at PO8. Error bars indicate the standard error. ns indicates a statistically nonsignificant difference, * indicates p < 0.05, * * indicates p < 0.01 and * * * indicates p < 0.001. The p values of all comparisons are shown in Table 2.

**Table 2.** Multiple comparison p values of N1 amplitudes at PO8 evoked by shapes in Experiment 2.

| Shape | PTS | UTS | SS | SC | QC |
| --- | --- | --- | --- | --- | --- |
| UTS | 0.0026 ** | — | — | — | — |
| SS | 0.0001 *** | 0.0005 *** | — | — | — |
| SC | 0.0002 *** | 0.0008 *** | 0.9759 ns | — | — |
| QC | 0.0001 *** | 0.1476 ns | 0.0159 * | 0.0237 * | — |
| KSI | 0.7438 ns | 0.0156 * | 0.0005 *** | 0.0005 *** | 0.0001 *** |

PTS, pyramidal triplet squares; UTS, unconnected triplet squares; SS, solitary square; SC, solitary circle; QC, quadruplet circles; KSI, Kanizsa square inducers.
ns, statistically nonsignificant difference; \* $p < 0.05$ ; \*\* $p < 0.01$ ; \*\*\* $p < 0.001$

**Fig. 7.**
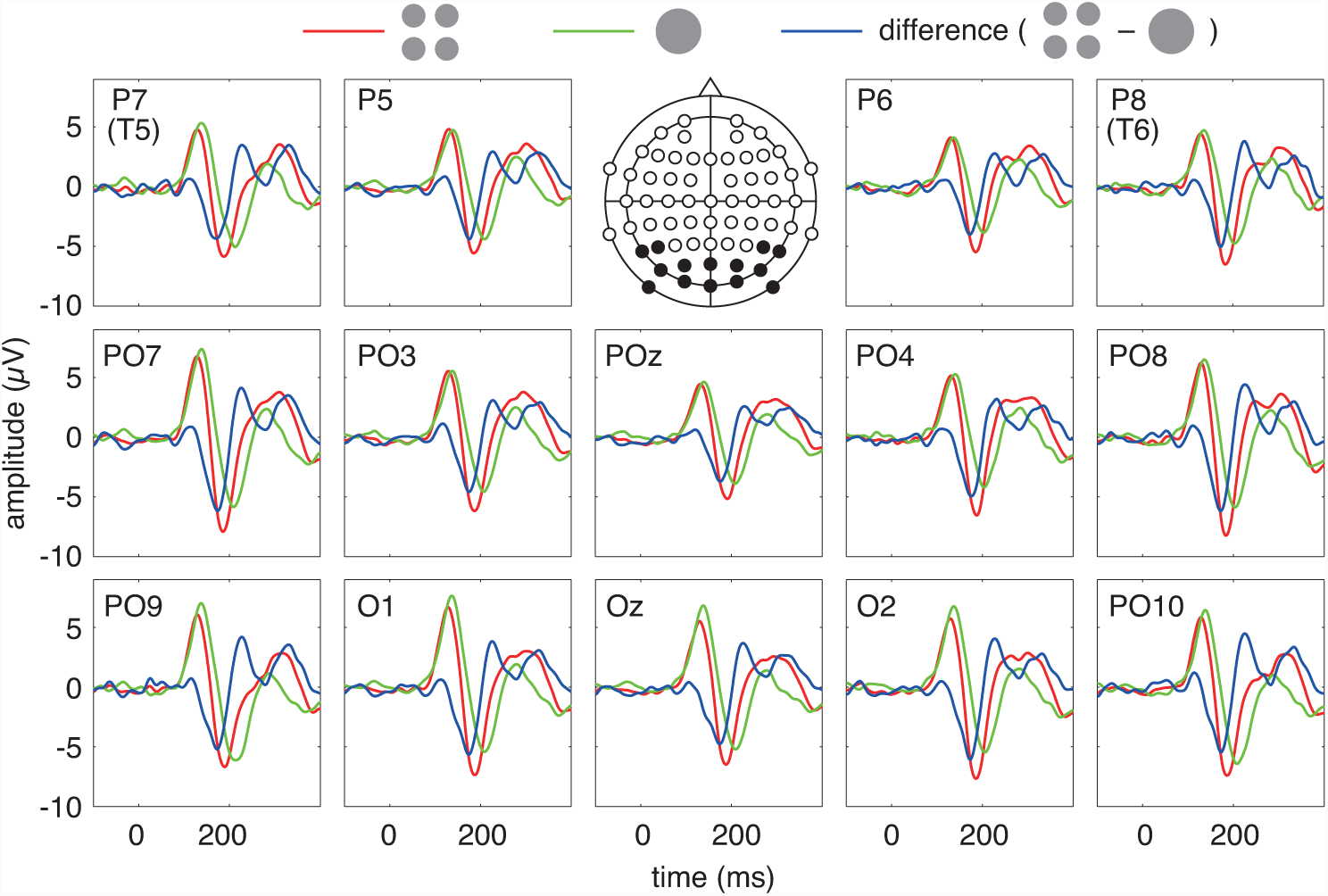
Grand-averaged (n = 16) ERPs evoked by quadruplet circles (QC; red line) and solitary circle (SC; green line) and the difference waves (QC – SC; blue line).

**Fig. 8.**
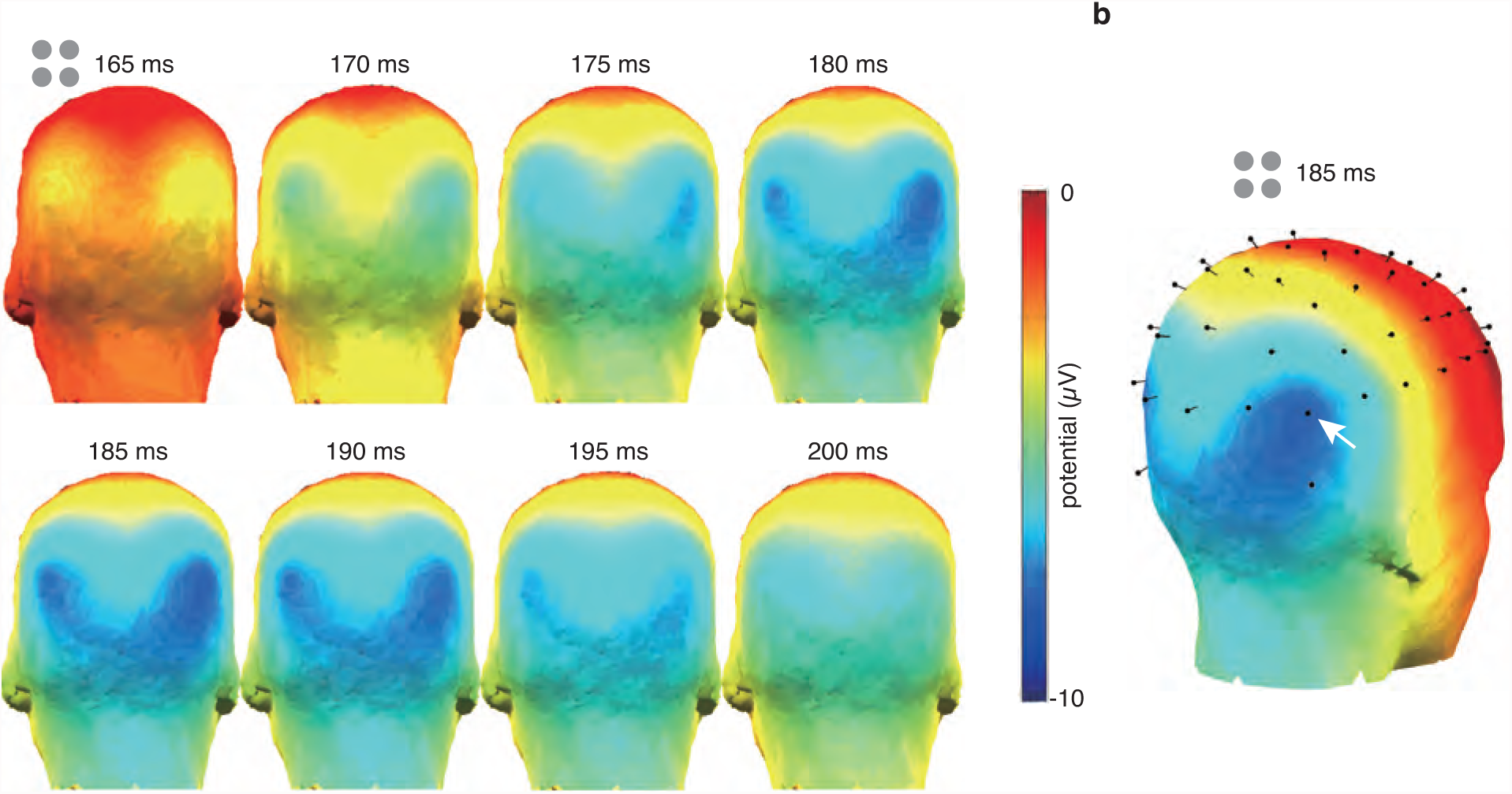
N1 scalp topography evoked by quadruplet circles (QC). (a) Chronological series from 160 ms to 195 ms. (b) Topography at the peak time (179 ms). The white arrow indicates the location of PO8.

Compared with TS and QC, SS and SC evoked a weaker N1 with approximately 20 ms slower peak latencies (SS, –6.7 ± 0.9 µV, 211 ± 4 ms; SC, –6.7 ± 1.0 µV, 203 ± 3 ms; Figs. 5-7). The N1 scalp topography by solitary shapes did not show sharp peaks in the anterior LO. They showed weakly activated areas that were broadly distributed in the occipital region (Fig. 9). Therefore, solitary shapes do not significantly or exclusively activate the anterior LO.

**Fig. 9.**
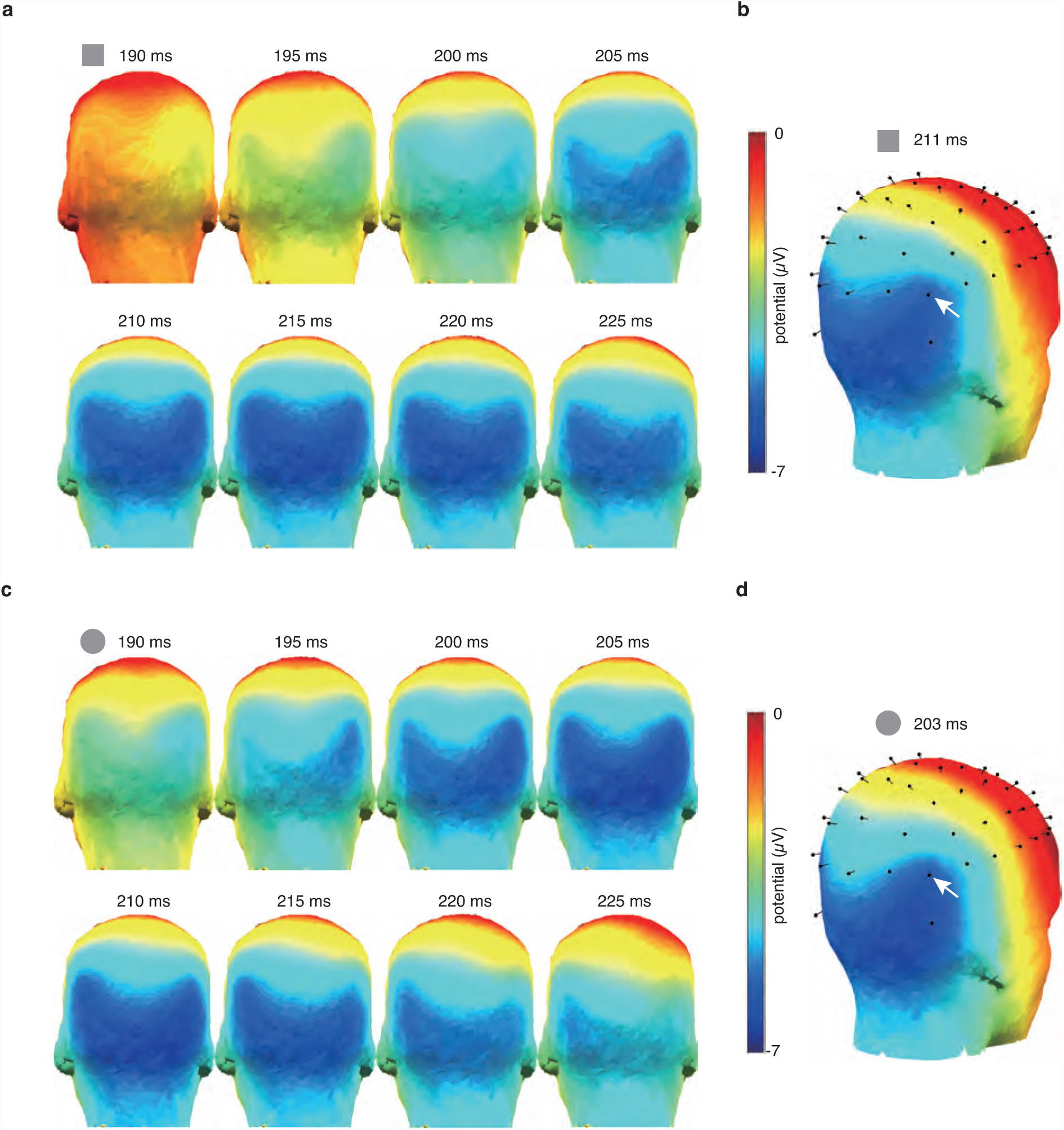
N1 scalp topography evoked by solitary shapes. (a) Solitary square (SS). Chronological series from 190 ms to 225 ms. (b) Solitary square (SS). Topography at the peak time (204 ms). (c) Solitary circle (SC). Chronological series from 190 ms to 225 ms. (d) Solitary circle (SC). Topography at the peak time (204 ms).

The difference waves between the ERPs evoked by PTS and SS and the different waves between the ERPs evoked by QC and SC are shown in Figs. 5 and 7, respectively. These data show that the negative peaks in the N1 time range were maximal at PO8 (square difference [PTS – SS], – 9.3 µV, 182 ms; circle difference [QC – SC], –6.2 µV, 172 ms). The scalp potential topographies of the difference waves showed sharp peaks in the anterior lateral occipital region (Fig. 10), suggesting that the multiplicity or compositeness of the shapes activates the anterior LO.

**Fig. 10.**
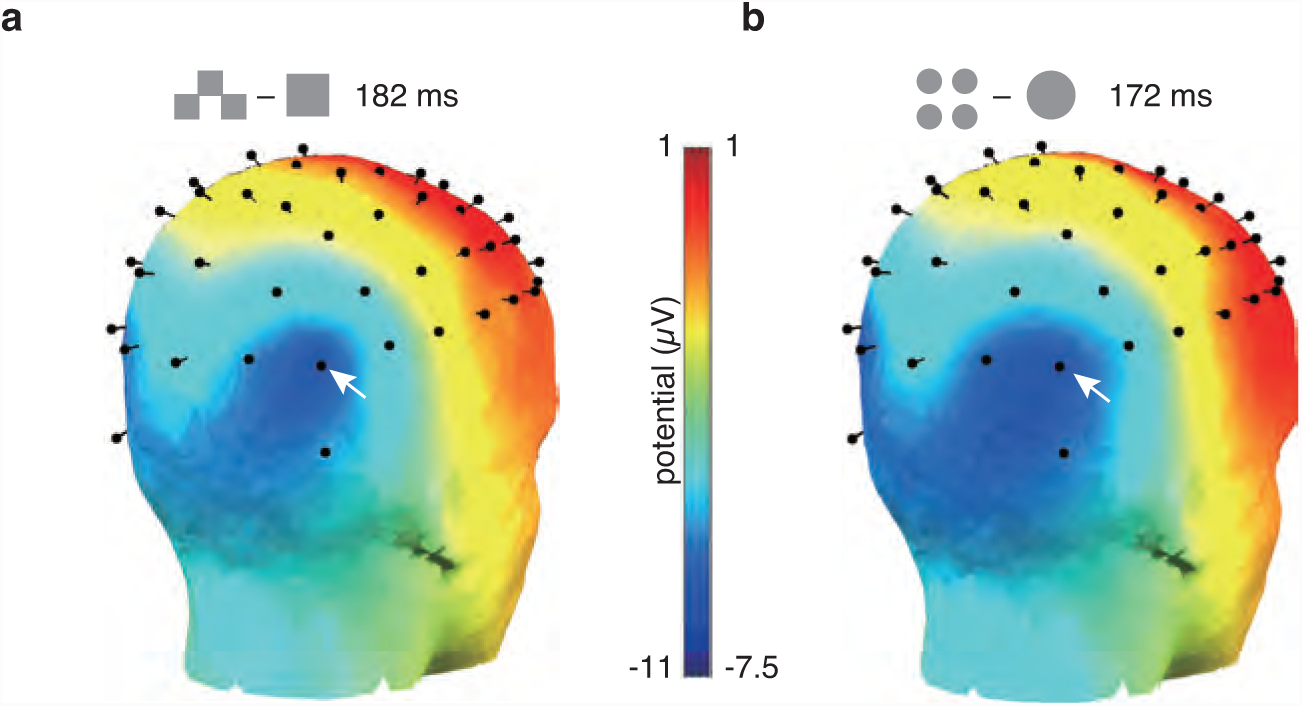
a) Scalp topography of difference waves between the ERPs evoked by pyramidal triplet squares (PTS) and solitary square (SS) at the peak time (182 ms). b) Scalp topography of difference waves between the ERPs evoked by quadruplet circles (QC) and solitary circle (SC) at the peak time (172 ms).

UTS evoked an 18% smaller N1 than PTS at PO8 (–10.5 ± 1.2 µV; paired t-test modified for multiple comparisons, p = 0.0026, Table 2; Figs. 6 and 11a), suggesting that the separation or unconnectedness of element shapes of composite figures decreases the activation of the LO.

KSI evoked an N1 component that was 38% larger at PO8 than that evoked by QC (–13.1 ± 1.6 µV; Figs. 6 and 11b). The amplitude of the N1 evoked by KSI was similar to the PTS response (the average of individual differences between N1 amplitudes evoked by KSI and PTS, 0.3 ± 0.8 µV; paired t-test modified for multiple comparisons, p = 0.7438, Table 2; Fig. 6). Thus, it has been confirmed that PTS evokes an N1 comparable to that evoked by KSI in the anterior LO. The difference waves between the ERPs evoked by PTS and UTS and the different waves between the ERPs evoked by KSI and QC are shown in Fig. 11. Negative peaks in the N1 time range were maximal at PO8.

**Fig. 11.**
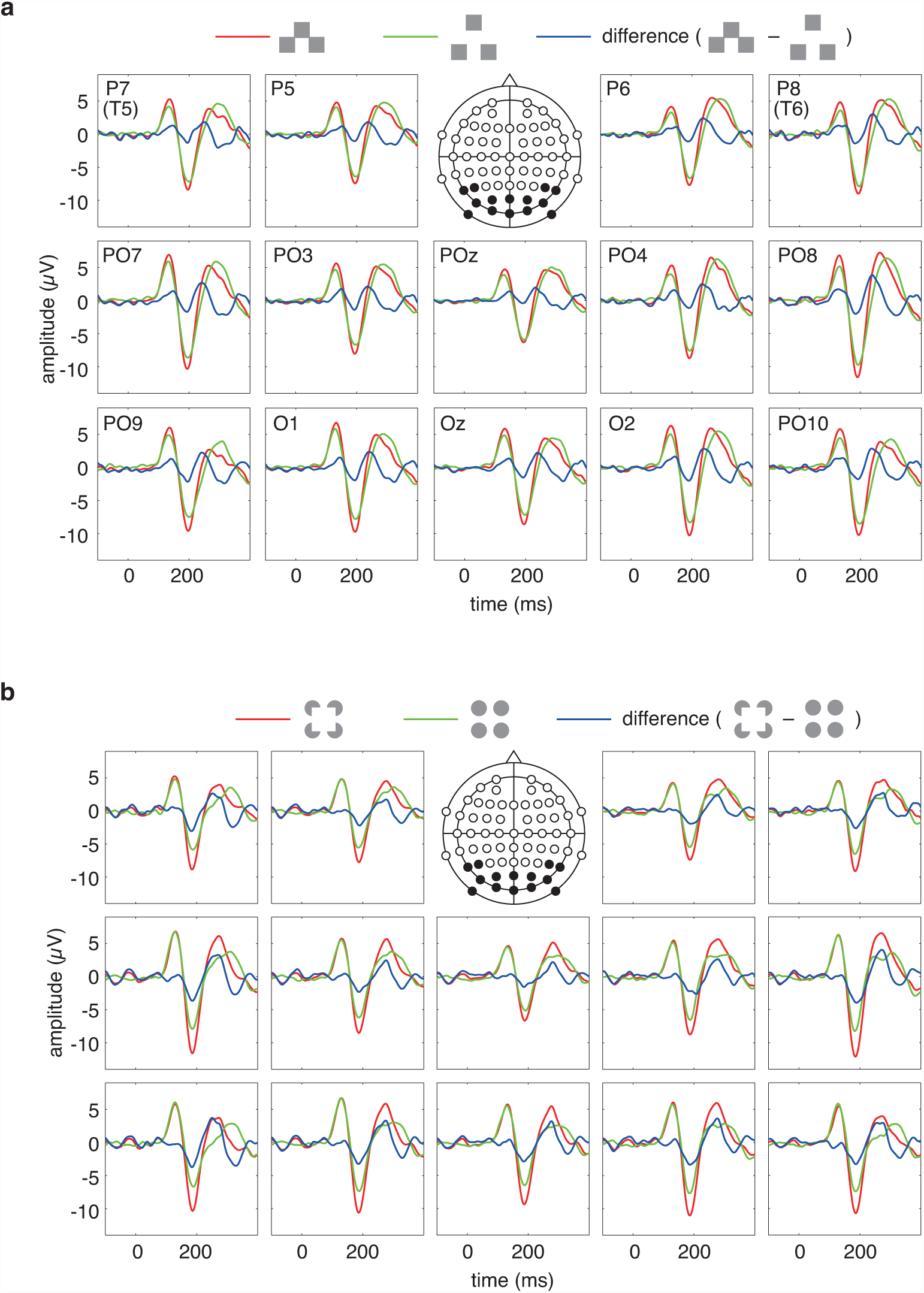
a) Grand-averaged (n = 16) ERPs evoked by pyramidal triplet squares (PTS; red line) and unconnected triplet squares (UTS; green line) and the difference waves (PTS – UTS; blue line). b) Grand-averaged (n = 16) ERPs evoked by Kanizsa square inducers (KSI; red line) and quadruplet circles (QC; green line) and the difference waves (KSI – QC; blue line).

The scalp potential topography of the N1 component evoked by UTS and KSI showed a similar pattern to those evoked by PTS and QC (Fig. 12), indicating that all of these multielement figures activate the same cortical area, the anterior LO. The scalp potential topographies of the difference waves (PTS – UTS and KSI – QC) showed sharp peaks in the anterior lateral occipital region (Fig. 13), suggesting that the compositeness or connectedness of shapes activates the anterior LO.

**Fig. 12.**
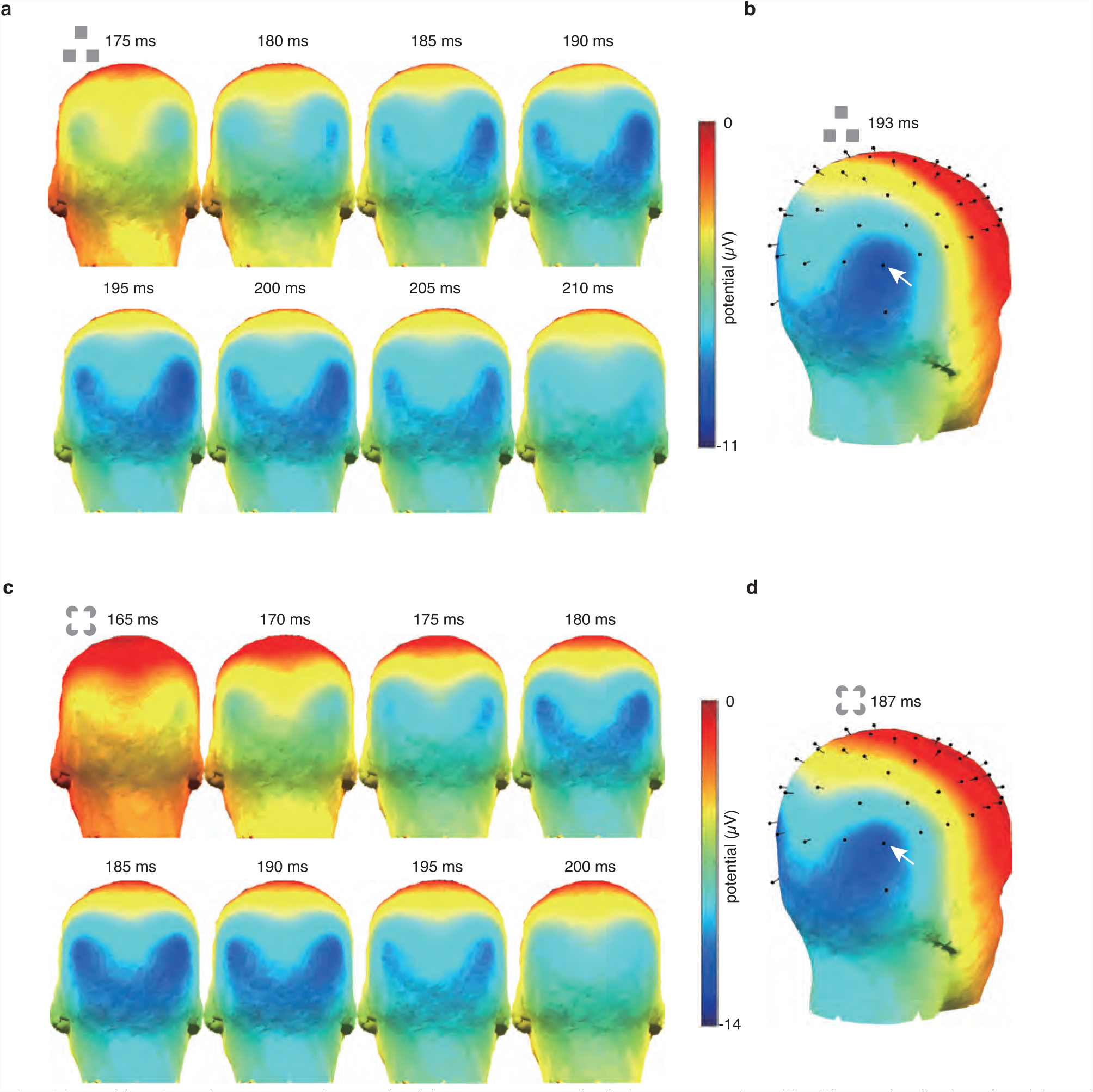
a, b) N1 scalp topography evoked by unconnected triplet squares (UTS). Chronological series (a) and topography at the peak time (b). c, d) N1 scalp topography evoked by Kanizsa square inducers (KSI). Chronological series (c) and topography at the peak time (d).

**Fig. 13.**
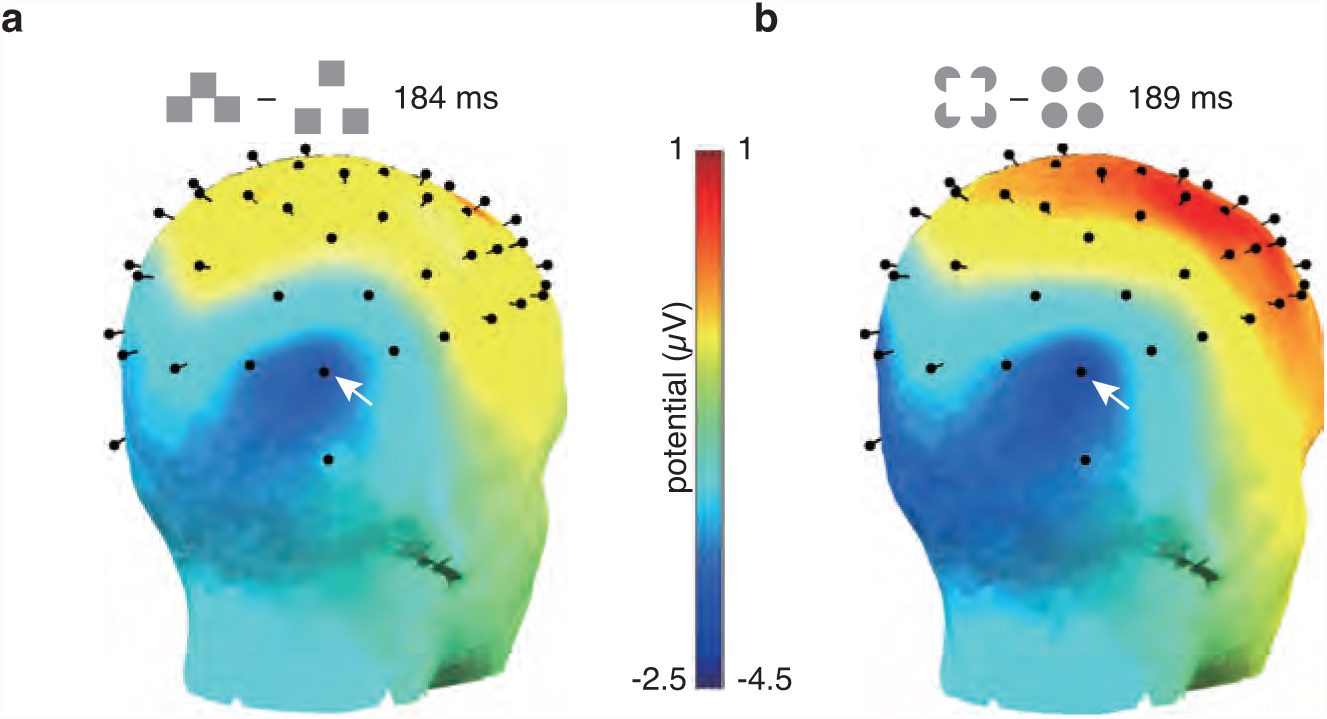
a) Scalp topography of difference waves between the ERPs evoked by pyramidal triplet squares (PTS) and unconnected TS (UTS) at the peak time (184 ms). b) Scalp topography of difference waves between the ERPs evoked by Kanizsa square inducers (KSI) and quadruplet circles (QC) at the peak time (189 ms).

### Experiment 3 (five figures composed of three squares)

We presented five figures composed of three squares: solitary long rectangle (LR, Fig. 1g), square and rectangle (SR, Fig. 1h), L-shape (LS, Fig. 1i), PTS (Fig. 1a) and oblique triplet squares (OTS, Fig. 1j). All shapes evoked a distinguishable N1 at PO8 across 15 participants. As in Experiments 1 and 2, four composite shapes (SR, LS, PTS and OTS) evoked a large N1 at the lateral occipital sites (Figs. 14, 15b-e and 16, Table 3). Compared with PTS, OTS also activated more medial regions (Fig. 15d-f). The N1 amplitude at PO8 by composite figures was significantly larger than that by solitary LR (Fig. 16, Table 3). There was no significant difference among the N1 amplitudes evoked by the four composite shapes.

**Fig. 14.**
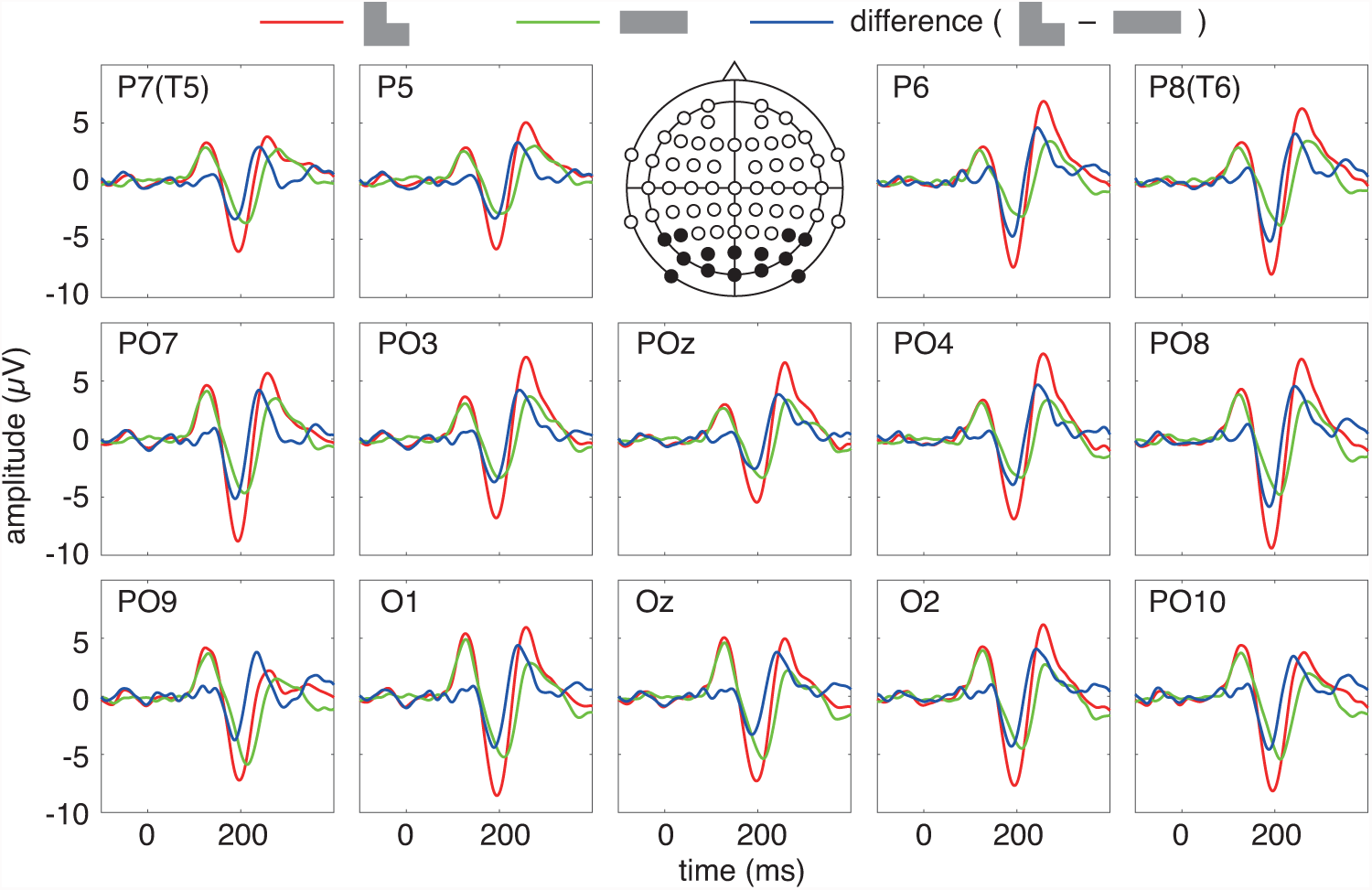
Grand-averaged (n = 15) ERPs evoked by L-shape (LS; red line) and long rectangle (LR; green line) and the difference waves (LS- LR; blue line).

**Fig. 15.**
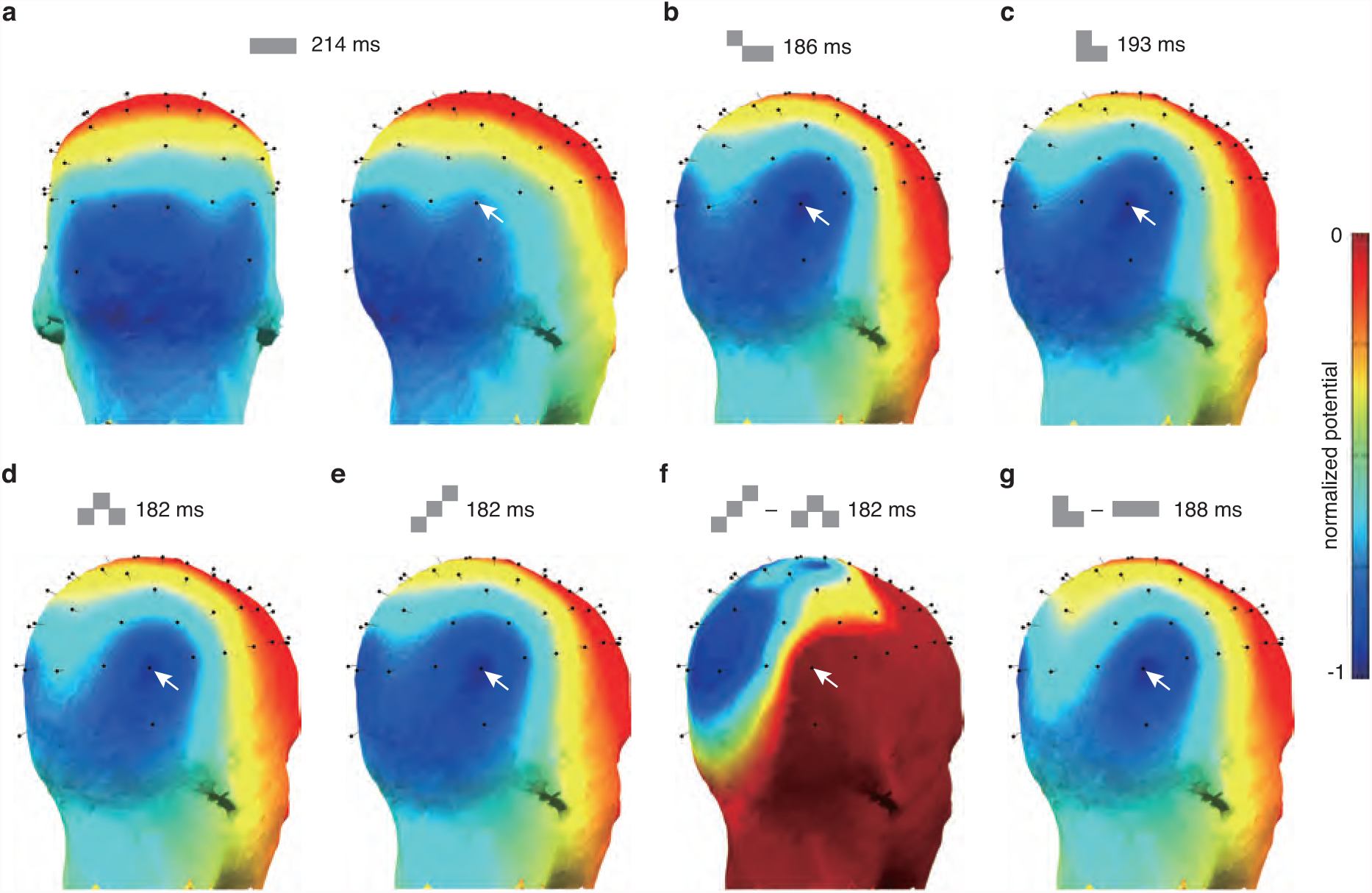
a-e) Normalized N1 scalp topographies evoked by long rectangle (LR, a), square and rectangle (SR, b), L-shape (LS, c), pyramidal triplet squares (PTS, d) and oblique triplet squares (OTS, e) at each peak time. f,g) Scalp topographies of normalized difference waves (f, OTS – PTS; g, LS – LR) at each peak time.

**Fig. 16.**
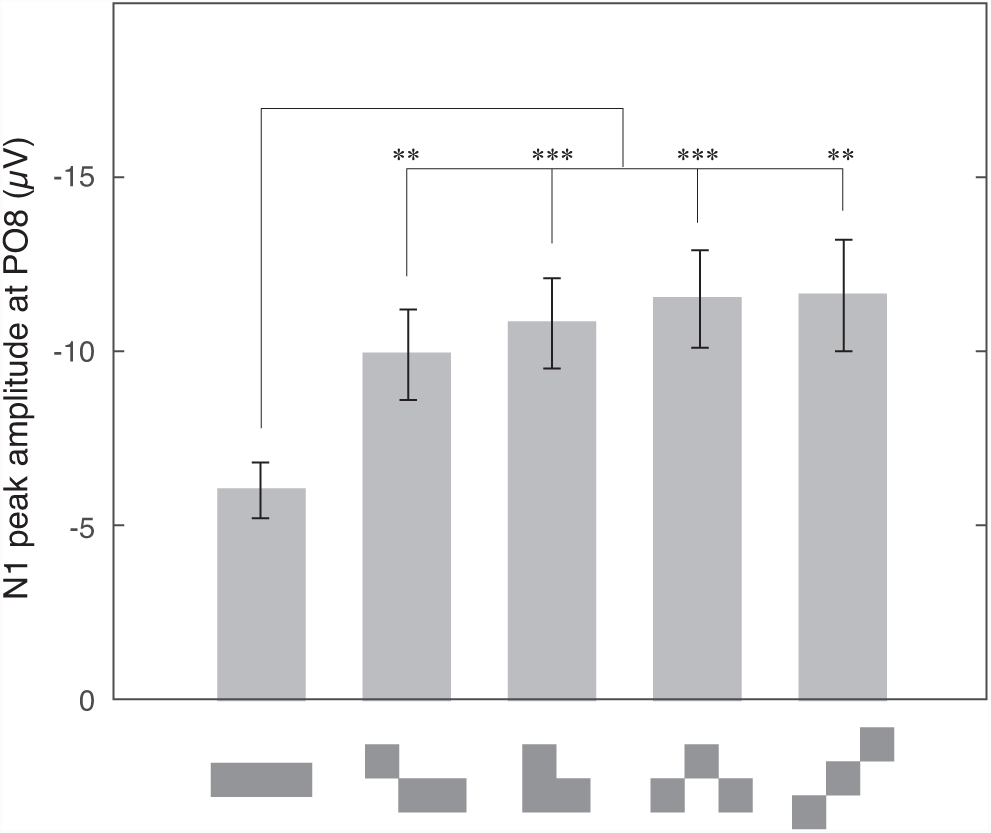
Averaged (n = 15) N1 peak amplitudes evoked by long rectangle (LR), square and rectangle (SR), L-shape (LS), pyramidal triplet squares (PTS) and oblique triplet squares (OTS) at PO8. Error bars indicate the standard error. * * indicates p < 0.01 and * * * indicates p < 0.001. The p values of all comparisons are shown in Table 3.

**Table 3.** Multiple comparison p values of N1 amplitudes at PO8 evoked by shapes in Experiment 3.

| Shape | LR | SR | LS | PTS |
| --- | --- | --- | --- | --- |
| SR | 0.0034 ** | — | — | — |
| LS | 0.0003 *** | 0.1126 | — | — |
| PTS | 0.0004 *** | 0.05 | 0.3314 | — |
| OTS | 0.0022 ** | 0.077 | 0.3581 | 0.9489 |

LR, long rectangle; SR, square and rectangle; LS, L-shape; PTS, pyramidal triplet squares; OTS, oblique triplet squares. \*\* $p < 0.01$ ; \*\*\* $p < 0.001$

Compared with the composite shapes, solitary LR evoked weaker N1 components with 20-30 ms slower peak latencies (Fig. 14), similar to SS and SC in Experiment 2. The N1 scalp topography by LR did not show sharp peaks in the anterior LO. They showed weakly activated areas that were broadly distributed in the occipital region (Fig. 15a). This scalp potential distribution is similar to that evoked by SS and SC in Experiment 2.

The difference waves between the ERPs evoked by LS and LR are shown in Fig. 14. These data show that the negative peaks in the N1 time range were maximal at PO8. The scalp potential topography of the difference waves showed sharp peaks in the anterior lateral occipital region (Fig. 15g).

## Discussion

### Shape analysis in the anterior LO

Malach and colleagues (1995) originally described the location of the lateral occipital complex (LOC) as “the lateral-posterior aspect of the occipital lobe, just abutting the posterior aspect of the motion-sensitive area MT/V5”. This original definition has been extended and revised based on results from recent imaging studies. To date, the LOC is thought to be divided into an anterior and ventral regions (the posterior fusiform gyrus or the ventral occipitotemporal cortex) and posterior and lateral regions, the LO (Grill-Spector & Malach, 2004; Kanwisher & Dilks, 2014; Wandell & Winawer, 2011). As shown in both previous and the present studies, the LO may be involved in shape analysis. Alternatively, the anterior and ventral regions of the LOC may be a part of the ventral temporal cortex (VTC), where the visual categorization of objects may take place (Grill-Spector & Weiner, 2014). Thus, the two regions of the LOC seem to differ in the hierarchy of the object recognition process. Because most studies prior to Larsson and Heeger (2006) and several recent LOC studies did not clearly distinguish between the two regions (rather, treat them as a unit), the results from these studies should be considered carefully, particularly for region-specific functional specialization.

Larsson and Heeger (2006) clarified that the LO is divided into two independent retinotopic areas (i.e., the posterior LO1 and anterior LO2) using fMRI. Although these authors confirmed that both LO1 and LO2 are “object-selective”, using the LOC functional localizer, based on enhanced responses to images of faces and 3D objects compared with responses to scrambled images of these stimuli, they also found that LO2 is significantly more responsive to objects than LO1. Furthermore, Larsson and colleagues found that LO1 exhibits orientation-selective responses to simple grating stimuli, whereas LO2 shows no selectivity for stimulus orientation (Larsson et al., 2006). This functional specialization within the LO was recently confirmed by a transcranial magnetic stimulation (TMS) study (Silson et al., 2013). Silson and colleagues (2013) performed fMRI-aided, localized TMS of LO1 and LO2 and found that the TMS of LO1 disrupted orientation, but not shape, discrimination, whereas the TMS of LO2 disrupted shape, but not orientation, discrimination. The present study also observed significant activity evoked by simple 2D composite figures composed of elementary shapes in the anterior LO, and BESA indicated that the source of the activation was localized at the LO2. Thus, the activity that we reported in this paper might reflect the neural processing of shape in the LO2.

Simple composite figures, such as PTS, may evoke significant activation of shape-analyzing processes in LO2, as shown in the present study. However, solitary elementary shapes barely activate the anterior LO. Thus, the anterior LO might not analyze the contour or surface of solitary elementary shapes. However, it might analyze the geometrical relationships among multiple 2D shapes that composed a single object. Similarly, fMRI studies by Murray and colleagues found that line drawings of simple 3D shapes composed of only straight lines that can be viewed as 2D composite figures significantly activated the LO compared with solitary 2D shapes (S. O. Murray, Kersten, Olshausen, Schrater, & Woods, 2002). Although a preliminary representation of surfaces oriented in depth has been considered necessary for the structural interpretation of 3D objects (Marr, 1982; Palmer, 1999), a recent alternative theory regarding the 3D perception of objects has been proposed (Pizlo, 2008; Pizlo, Sawada, Li, Kropatsch, & Steinman, 2010). The theory claims that if several a priori constraints representing the properties of 3D shapes (including maximal planarity of contour) are applied, the 3D shape of an object may subsequently be reconstructed from only one of its 2D images. For the simplest example, a line drawing of an opaque cube composed of three tetragons (e.g., one top rhombus and two side parallelograms) is vividly perceived as a 3D cube. In contrast, the PTS used in the present study that is similarly composed of three squares does not evoke a 3D perception. Thus, the possible analysis of geometrical relationships among element shapes of composite figures implemented in the anterior LO might be essential for structural interpretations of 2D and 3D objects. Only a tiny fraction of 2D composite figures was examined in the present study. Therefore, more systematic and comprehensive studies that present various 2D/3D composite figures must be performed,.

Sugawara and Morotomi first reported that IC figures, including KSI, evoked a significant N1 in the lateral occipital region (Sugawara & Morotomi, 1991). Their recording sites were 4 cm above the inion and 5 cm to the left and right of the midline, which are similar locations to PO7 and PO8 in the present study. To date, many studies, including fMRI studies, have investigated the activation in the LO evoked by IC figures (see (M. M. Murray & Herrmann, 2013) for a review). These studies have revealed that Kanizsa-type inducers alone evoke a significant N1 in the LO, even when the inducers are oriented to not form an IC. Moreover, these inducers may evoke a much larger N1 in the LO if they are oriented to form an IC. We showed that the QC (which are circles and not truly KSI) evoked a large N1 in the LO. However, KSI evoked a larger N1 in the anterior LO than QC. If KSI are properly oriented, participants perceive an illusory square and four circles partially occluded by the illusory square. Partial occlusion is an important cue for detecting the geometrical relationships between element shapes of composite figures (Palmer, 1999). Thus, occluded shapes appear as far from and occluding shapes appear as near to the viewer. Therefore, the analysis of geometrical relationships among shapes that compose an object likely includes an inspection of partial occlusion between shapes.

As Gestalian psychologists have demonstrated, certain visual elements are instantly and effortlessly organized into objects depending on several of their geometric properties (Kanizsa, 1979; Palmer, 1999; Wagemans, Elder, et al., 2012a; Wagemans, Feldman, et al., 2012b). Although the neural mechanisms for this perceptual organization remain largely unknown (Wagemans, Elder, et al., 2012a; Wagemans, Feldman, et al., 2012b), some perceptual organization mechanisms have been hypothesized to be implemented in the LO or LOC (see (Sasaki, 2007) for a review of early studies). For a recent example, based on their TMS experiments, Bona and colleagues claim that the perceptual grouping of Gabor patches into contours occur in the LO (Bona, Herbert, Toneatto, Silvanto, & Cattaneo, 2014). Similarly the present results may be interpreted according to the context of perceptual organization. Thus, the N1 evoked by simple multielement figures in the anterior LO might reflect the activity of the perceptual grouping processes implemented in the anterior LO. Kim and Biederman found that line drawings of two 3D objects further activate the LO if they are in contact or overlapping compared with objects that are separated (Kim & Biederman, 2012). The present study also indicated that physical and perceived composite figures such as PTS and KSI evoke a larger N1 in the anterior LO than unconnected multielement figures. Palmer and Rock hypothesized that perceptual grouping and parsing for parts analysis may occur in parallel after processing figure-ground segmentation (Palmer & Rock, 1994). The parsing process for physical and perceived composite shapes might evoke an additional activation of the shape-analyzing process in the anterior LO. Symmetry is also an important attribute of figural goodness, and experimental evidence indicates that figure symmetry is processed in the LO (Bona et al., 2014; Sasaki, Vanduffel, Knutsen, Tyler, & Tootell, 2005), supporting the hypothesis that the LO participates in perceptual organization.

### N1/N170 evoked by shapes and faces

The definition of N170 varies slightly among ERP experts. Rossion and Jacques believe that N170 is identical to the visual N1 (Rossion & Jacques, 2011); however, Luck regards the N170 as a subcomponent of the N1 (Luck, 2014). Referred to as the “N170 face effect”, it is widely believed that the largest N1/N170 response is evoked by faces (Rossion, 2014; Rossion & Jacques, 2008; 2011). The present study showed that simple composite shapes such as PTS and KSI could evoke significant N1 amplitudes comparable with the amplitudes evoked by faces. However, we found that the cortical areas activated by faces and simple composite figures differ. The sources of N1 evoked by PTS were localized in the anterior LO in the present study, whereas dominant sources of the N1/N170 evoked by faces were localized in the lateral portion of the posterior fusiform gyrus and in the anterior/middle fusiform gyrus (Rossion & Jacques, 2011).

Although Malach and colleagues (1995) reported that all three categories objects (faces, familiar objects and unfamiliar abstract sculptures) evenly activate the LOC compared with images of textures, Larsson and Heeger (2006) found that both the LO1 and LO2 responded more strongly to images of objects than to images of faces. The present study also found that TS evoked 24% larger responses at PO8 than faces. Moreover, previous cortical surface recordings have revealed that various simple figures evoke a large (up to more than 100 µV) N1 locally at the LO (Allison, Puce, Spencer, & McCarthy, 1999; Fiebelkorn, Foxe, Schwartz, & Molholm, 2010; Sehatpour et al., 2008). Because human faces are also complex 3D objects, the geometrical complexity of objects alone is unlikely to be reflected in the activation of LO2.

Judging from the present results, artificially manipulated faces might activate both face and shape processing areas. For example, the discontinuity of misaligned top and bottom halves of faces in the method introduced by Young and colleagues (Young, Hellawell, & Hay, 1987) might induce the perception of two misaligned planes textured with faces, and the perceived composite figures might also activate the anterior LO. In fact, Letourneau and Mitchell found that misaligned faces evoked significant N1/N170 at the TO2 electrode (which is geometrically comparable with the PO8 electrode in the present study), but intact faces did not (Letourneau & Mitchell, 2008).

Recently, Thierry and colleagues challenged the “N170 face effect” by claiming that the control of interstimulus perceptual variance (ISPV) abolishes N1/N170 face selectivity. They also claimed that matched ISPV cars evoked larger N1/N170 amplitudes than faces (Dering, Martin, & Thierry, 2009; Thierry, Martin, Downing, & Pegna, 2007). However, their claims evoked prompt criticism from international leaders in the field (Bentin et al., 2007). In particular, Rossion and Jacques examined the data by Thierry and colleagues and thoroughly criticized them in their review paper (Rossion & Jacques, 2008). Because our study was not designed to explain this discrepancy, we do not have direct evidence to settle the controversy. However, if a clue regarding this discrepancy exists in the present study, it might be that the configuration attributes of cars should be carefully examined. In particular, the front views of cars might have composite geometric features and caricaturized face-like features; therefore, this view might activate both the face areas and the anterior LO, in addition to other higher visual areas. Therefore, because the activation amplitudes might depend on a combination of the various configuration properties of those stimuli, pictures of cars might evoke larger N1/N170 amplitudes than faces in certain experiments. In addition, the selection of electrodes for averaging might significantly affect N1/N170 amplitudes because the scalp topographies of N1 differ in response to shapes and faces, as shown in the present study.

## Acknowledgments

We thank Dr. Y. Sasaki for the critical comments on an early draft of this paper. We also thank former students N. Nishi, K. Mitsuishi, Y. Kubota and K. Sueto for their assistance with the ERP recordings during the preliminary stage of this project.

---

**Fig. S1.**
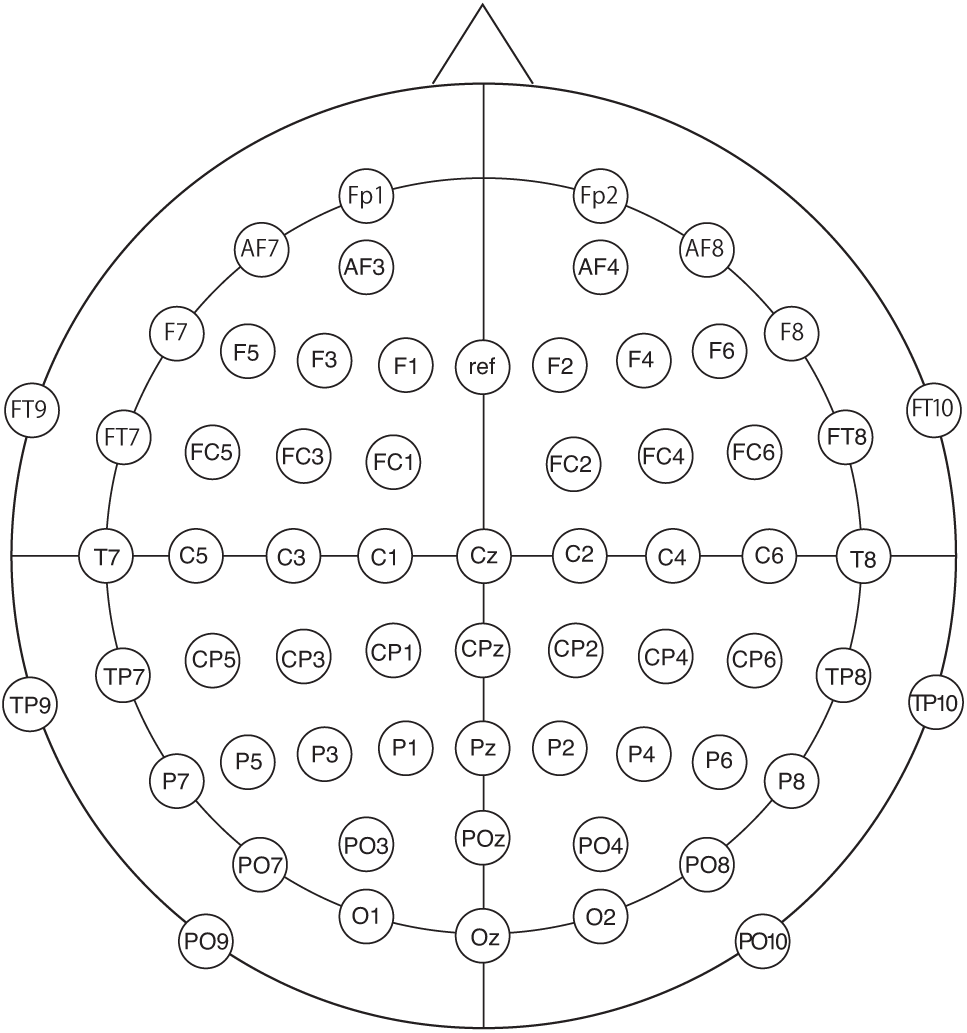
Electrode placement in the present study. Sixty-four electrodes were placed according to the international 10-10 system.

**Fig. S2.**
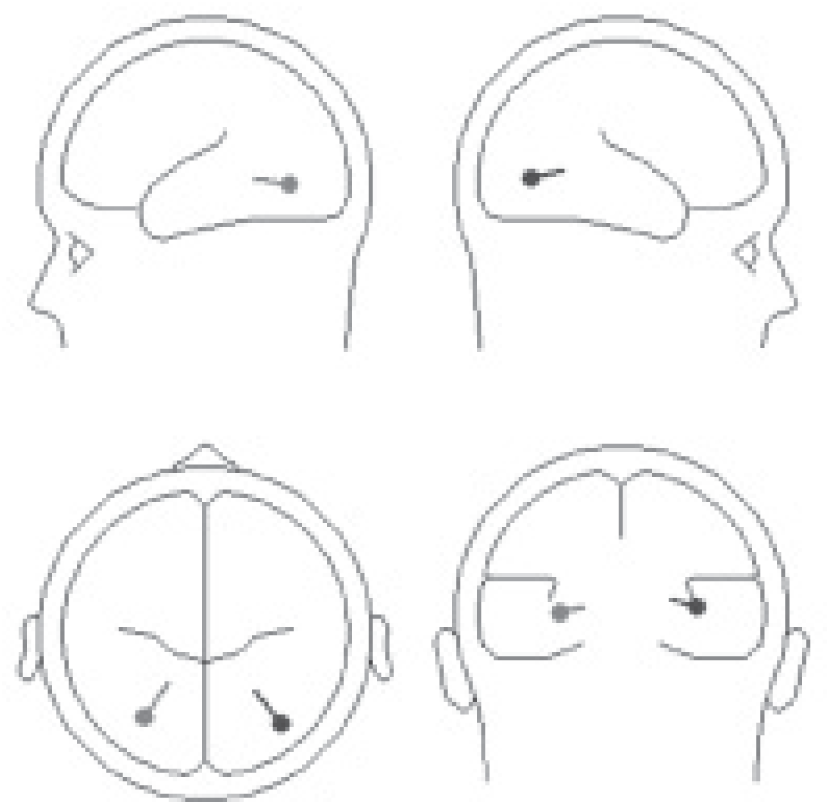
Current dipoles of N1 evoked by PTS that were estimated using BESA.

## References

Allison, T., Puce, A., Spencer, D. D., & McCarthy, G. (1999). Electrophysiological studies of human face perception. I: Potentials generated in occipitotemporal cortex by face and non-face stimuli. Cerebral Cortex (New York, N.Y.: 1991), 9(5), 415–430.

Bentin, S., Taylor, M. J., Rousselet, G. A., Itier, R. J., Caldara, R., Schyns, P. G., et al. (2007). Controlling interstimulus perceptual variance does not abolish N170 face sensitivity. Nature Neuroscience, 10(7), 801–2– author reply 802–3. http://doi.org/10.1038/nn0707-801

Bona, S., Herbert, A., Toneatto, C., Silvanto, J., & Cattaneo, Z. (2014). The causal role of the lateral occipital complex in visual mirror symmetry detection and grouping: an fMRI-guided TMS study. Cortex, 51, 46–55. http://doi.org/10.1016/j.cortex.2013.11.004

Delorme, A., & Makeig, S. (2004). EEGLAB: an open source toolbox for analysis of single-trial EEG dynamics including independent component analysis. Journal of Neuroscience Methods, 134(1), 9–21. http://doi.org/10.1016/j.jneumeth.2003.10.009

Dering, B., Martin, C. D., & Thierry, G. (2009). Is the N170 peak of visual event-related brain potentials car-selective? NeuroReport, 20(10), 902–906. http://doi.org/10.1097/WNR.0b013e328327201d

Fiebelkorn, I. C., Foxe, J. J., Schwartz, T. H., & Molholm, S. (2010). Staying within the lines: the formation of visuospatial boundaries influences multisensory feature integration. The European Journal of Neuroscience, 31(10), 1737–1743. http://doi.org/10.1111/j.1460-9568.2010.07196.x

Grill-Spector, K., & Malach, R. (2004). The human visual cortex. Annual Review of Neuroscience, 27, 649–677. http://doi.org/10.1146/annurev.neuro.27.070203.144220

Grill-Spector, K., & Weiner, K. S. (2014). The functional architecture of the ventral temporal cortex and its role in categorization. Nature Reviews Neuroscience, 15(8), 536–548. http://doi.org/10.1038/nrn3747

Hatta, T., & Nakatsuka, Z. (1976). Note on hand preference of Japanese people. Perceptual and Motor Skills, 42(2), 530–530. http://doi.org/10.2466/pms.1976.42.2.530

Kanizsa, G. (1979). Organization in vision. Praeger Publishers.

Kanwisher, N., & Dilks, D. D. (2014). The Functional Organization of the Ventral Visual Pathway in Humans. In L. M. Chalupa & J. Werner (Eds.), The New Visual Neurosciences.

Kim, J. G., & Biederman, I. (2012). Greater sensitivity to nonaccidental than metric changes in the relations between simple shapes in the lateral occipital cortex. NeuroImage, 63(4), 1818–1826. http://doi.org/10.1016/j.neuroimage.2012.08.066

Larsson, J., & Heeger, D. J. (2006). Two retinotopic visual areas in human lateral occipital cortex. The Journal of Neuroscience: the Official Journal of the Society for Neuroscience, 26(51), 13128–13142. http://doi.org/10.1523/JNEUROSCI.1657-06.2006

Larsson, J., Landy, M. S., & Heeger, D. J. (2006). Orientation-selective adaptation to first- and second-order patterns in human visual cortex. Journal of Neurophysiology, 95(2), 862–881. http://doi.org/10.1152/jn.00668.2005

Letourneau, S. M., & Mitchell, T. V. (2008). Behavioral and ERP measures of holistic face processing in a composite task. Brain and Cognition, 67(2), 234–245. http://doi.org/10.1016/j.bandc.2008.01.007

Lopez-Calderon, J., & Luck, S. J. (2014). ERPLAB: an open-source toolbox for the analysis of event-related potentials. Frontiers in Human Neuroscience, 8, 213. http://doi.org/10.3389/fnhum.2014.00213

Luck, S. J. (2014). An Introduction to the Event-Related Potential Technique (Second). MIT Press.

Malach, R., Reppas, J. B., Benson, R. R., Kwong, K. K., Jiang, H., Kennedy, W. A., et al. (1995). Object-related activity revealed by functional magnetic resonance imaging in human occipital cortex. Proceedings of the National Academy of Sciences, 92(18), 8135–8139.

Marr, D. (1982). Vision. New York: W.H. Freeman & Co.

Murray, M. M., & Herrmann, C. S. (2013). Illusory contours: a window onto the neurophysiology of constructing perception. Trends in Cognitive Sciences, 17(9), 471–481. http://doi.org/10.1016/j.tics.2013.07.004

Murray, S. O., Kersten, D., Olshausen, B. A., Schrater, P., & Woods, D. L. (2002). Shape perception reduces activity in human primary visual cortex. Proceedings of the National Academy of Sciences, 99(23), 15164–15169. http://doi.org/10.1073/pnas.192579399

Palmer, S. E. (1999). Vision Science. Bradford Books.

Palmer, S., & Rock, I. (1994). Rethinking perceptual organization: The role of uniform connectedness. Psychonomic Bulletin & Review, 1(1), 29–55. http://doi.org/10.3758/BF03200760

Pizlo, Z. (2008). 3D Shape. MIT Press.

Pizlo, Z., Sawada, T., Li, Y., Kropatsch, W. G., & Steinman, R. M. (2010). New approach to the perception of 3D shape based on veridicality, complexity, symmetry and volume. Vision Research, 50(1), 1–11. http://doi.org/10.1016/j.visres.2009.09.024

Rossion, B. (2014). Understanding face perception by means of human electrophysiology. Trends in Cognitive Sciences, 18(6), 310–318. http://doi.org/10.1016/j.tics.2014.02.013

Rossion, B., & Jacques, C. (2008). Does physical interstimulus variance account for early electrophysiological face sensitive responses in the human brain? Ten lessons on the N170. NeuroImage, 39(4), 1959–1979. http://doi.org/10.1016/j.neuroimage.2007.10.011

Rossion, B., & Jacques, C. (2011). The N170: Understanding the Time Course of Face Perception in the Human Brain. In E. S. Kappenman & S. J. Luck (Eds.), The Oxford Handbook of Event-Related Potential Components (pp. 115–141). Oxford University Press.

Sasaki, Y. (2007). Processing local signals into global patterns. Current Opinion in Neurobiology, 17(2), 132–139. http://doi.org/10.1016/j.conb.2007.03.003

Sasaki, Y., Vanduffel, W., Knutsen, T., Tyler, C., & Tootell, R. (2005). Symmetry activates extrastriate visual cortex in human and nonhuman primates. Proceedings of the National Academy of Sciences, 102(8), 3159–3163. http://doi.org/10.1073/pnas.0500319102

Sehatpour, P., Molholm, S., Schwartz, T. H., Mahoney, J. R., Mehta, A. D., Javitt, D. C., et al. (2008). A human intracranial study of long-range oscillatory coherence across a frontaloccipital-hippocampal brain network during visual object processing. Proceedings of the National Academy of Sciences, 105(11), 4399–4404. http://doi.org/10.1073/pnas.0708418105

Silson, E. H., McKeefry, D. J., Rodgers, J., Gouws, A. D., Hymers, M., & Morland, A. B. (2013). Specialized and independent processing of orientation and shape in visual field maps LO1 and LO2. Nature Neuroscience, 16(3), 267–269. http://doi.org/10.1038/nn.3327

Sugawara, M., & Morotomi, T. (1991). Visual evoked potentials elicited by subjective contour figures. Scandinavian Journal of Psychology, 32(4), 352–357.

Thierry, G., Martin, C. D., Downing, P., & Pegna, A. J. (2007). Controlling for interstimulus perceptual variance abolishes N170 face selectivity. Nature Neuroscience, 10(4), 505–511. http://doi.org/10.1038/nn1864

Thomaz, C. E., & Giraldi, G. A. (2010). A new ranking method for principal components analysis and its application to face image analysis. Image and Vision Computing, 28(6), 902–913. http://doi.org/10.1016/j.imavis.2009.11.005

Uchiyama, H., Mitsuishi, K., & Ohno, H. (2009). Random Walker Test: a computerized alternative to the Road-Map Test. Behavior Research Methods, 41(4), 1242–1253. http://doi.org/10.3758/BRM.41.4.1242

Vogel, E. K., & Luck, S. J. (2000). The visual N1 component as an index of a discrimination process. Psychophysiology, 37(2), 190–203.

Wagemans, J., Elder, J. H., Kubovy, M., Palmer, S. E., Peterson, M. A., Singh, M., & Heydt von der, R. (2012a). A century of Gestalt psychology in visual perception: I. Perceptual grouping and figure-ground organization. Psychological Bulletin, 138(6), 1172–1217. http://doi.org/10.1037/a0029333

Wagemans, J., Feldman, J., Gepshtein, S., Kimchi, R., Pomerantz, J. R., van der Helm, P. A., & van Leeuwen, C. (2012b). A century of Gestalt psychology in visual perception: II. Conceptual and theoretical foundations. Psychological Bulletin, 138(6), 1218–1252. http://doi.org/10.1037/a0029334

Wandell, B. A., & Winawer, J. (2011). Imaging retinotopic maps in the human brain. Vision Research, 51(7), 718–737. http://doi.org/10.1016/j.visres.2010.08.004

Young, A. W., Hellawell, D., & Hay, D. C. (1987). Configurational information in face perception. Perception, 16(6), 747–759.

